# GLOBAL ANALYSIS OF PROTRUSION-TRANCSRIPTOMES IDENTIFIES DISTINCT CATEGORIES OF LOCALIZED RNAS IN NON-NEURONAL CELLS

**DOI:** 10.64898/2026.09.17.752430

**Authors:** Briana Hojo, Shikha Kumari, Dominic Milla, Victoria Vang, Megan L Norris

## Abstract

RNA localization to protrusions in non-neuronal cells is an emerging molecular process, distinct from the canonical mechanism characterized in neurons. Here, we describe a fractionation workflow optimized for non-neuronal cells that increases sensitivity and reproducibility of transcriptome-wide quantification of protrusion-localized RNAs. Using the optimized method, we identify six categories of protrusion-localized RNAs in non-neuronal cells from mice, including long non-coding RNAs and pseudogenes, and observe spatially distinct subsets of mitochondrially-localized RNAs. Taken together, our results reveal previously unappreciated spatial regulation of diverse transcripts and point toward broadly acting unifying principles that extend across cell types and species.

## INTRODUCTION

Subcellular localization of RNAs is a pervasive, highly conserved process with diverse physiological roles in vertebrates. In neurons, mRNAs localized in protrusions (axons and dendrites) are locally translated to generate localized proteins, which is essential for learning and memory (reviewed in (Engel et al., 2020; Zappulo et al., 2017)). This process of localized mRNA/localized translation/localized protein can be considered the “canonical” mRNA localization mechanism. mRNAs also localize to protrusions of non-neuronal mesenchymal cells such as embryonic fibroblasts and metastatic cancer cells, and protrusion-localized mRNAs in non-neuronal cells have been shown to play a role in tissue morphogenesis and cell behavior *in vivo* (Costa et al., 2020; Dermit et al., 2020; Mason et al., 2025; Norris & Mendell, 2023; Wang et al., 2017). However, global mRNA localization to protrusions in non-neuronal cell types does not correlate with protein localization to protrusions (Mardakheh et al., 2015). This lack of correlation suggests there is an alternative or “non-canonical” function for mRNA localization to protrusions in non-neuronal cells. As such, the regulatory and functional mechanisms of RNA localization in non-neuronal cells cannot be directly inferred from neurons and must instead be empirically characterized.

Recent breakthroughs have begun this effort by uncovering a role for mRNA localization in regulating protein binding partner behavior. Specifically, changing mRNA localization changes how the encoded protein interacts with other proteins. So far, this mechanism has been shown to affect four protrusion-localized mRNAs (*Kif1c, Net1, Rab13,* and *Trak2)*(Bradbury et al., 2025; Gasparski et al., 2023; Moissoglu et al., 2020; Norris & Mendell, 2023). Strikingly, all four of these mRNAs require the same *trans-*factors and *cis*-regulatory elements for localization: they rely on the kinesin motor protein KIF1C and the tumor suppressor APC for trafficking, and have similar GA-rich *cis-*localization elements in their 3’UTRs (Chrisafis et al., 2020; Moissoglu et al., 2020; Norris & Mendell, 2023; Pichon et al., 2021; Wang et al., 2017). These data suggest that at least some portion of protrusion-localized mRNAs use shared regulatory networks to achieve a common functional outcome. Consistent with this idea, another striking subset of co-regulated protrusion-localized mRNAs are the ribosomal protein mRNAs (Dermit et al., 2020; Fusco et al., 2021; Shigeoka et al., 2019; Wang et al., 2017). Although the function of this peculiar localization pattern is not understood, localization of ribosomal protein mRNAs to cell protrusions is conserved in humans and known to require a common *trans*-factor, LARP1, and a common *cis*-localization element in the 5’UTR called a TOP motif (Dermit et al., 2020; Goering et al., 2023). Thus, ribosomal protein mRNAs represent a second subset of co-regulated protrusion-localized mRNAs in non-neuronal cells.

When viewed together, the subsets described above, the four APC/KIF1C-dependent mRNAs and ∼79 ribosomal protein mRNAs, represent at most 83 of the multiple hundreds of RNAs that localize to protrusions in non-neuronal cells. Intriguingly, other mRNAs that are not as well characterized have been reported to require KIF1C and/or APC for localization to protrusions, suggesting that additional mRNAs may fall within the APC/KIF1C-dependent subset (Pichon et al., 2021; Wang et al., 2017). Even with these potential additions, there remains a substantial blind spot in our understanding of how most protrusion-localized RNAs are regulated and what functional purpose they serve. At the same time, the co-regulatory nature of the APC/KIF1C-dependent mRNAs and the ribosomal protein mRNAs suggests there may be unifying principles that work across large subsets of protrusion-localized RNAs, and that by identifying these subsets and their shared principles, we can decrease the complexity of the system and make more rapid progress toward understanding RNA localization to protrusions as a whole.

Our goal to identify unifying principles and regulatory networks relies on the ability to conduct robust, unbiased, transcriptome-wide analyses of protrusion-localized RNAs. However, persistent methodological issues, described below, have so far largely constrained these types of studies in non-neuronal cell types. The gold standard for high-throughput, transcriptome-wide analysis of protrusion-localized RNAs in neurons is the Transwell fractionation method (Taliaferro, 2019). In this method, cells are plated on membranes with microscopic pores, which protrusions, but not cell bodies, can pass through. Cell bodies are scraped from the top of the membrane and protrusions are lysed from within and beneath the membrane (Figure 1A). To date, application of this method in non-neuronal mesenchymal cells, which are smaller and more dynamic than neurons, has suffered from poor reproducibility across experiments and low sensitivity for detecting differences between localized and diffuse RNAs, making it difficult to distinguish bona fide biological differences from technical noise (Norris & Mendell, 2023; Wang et al., 2017).

**Figure 1.**
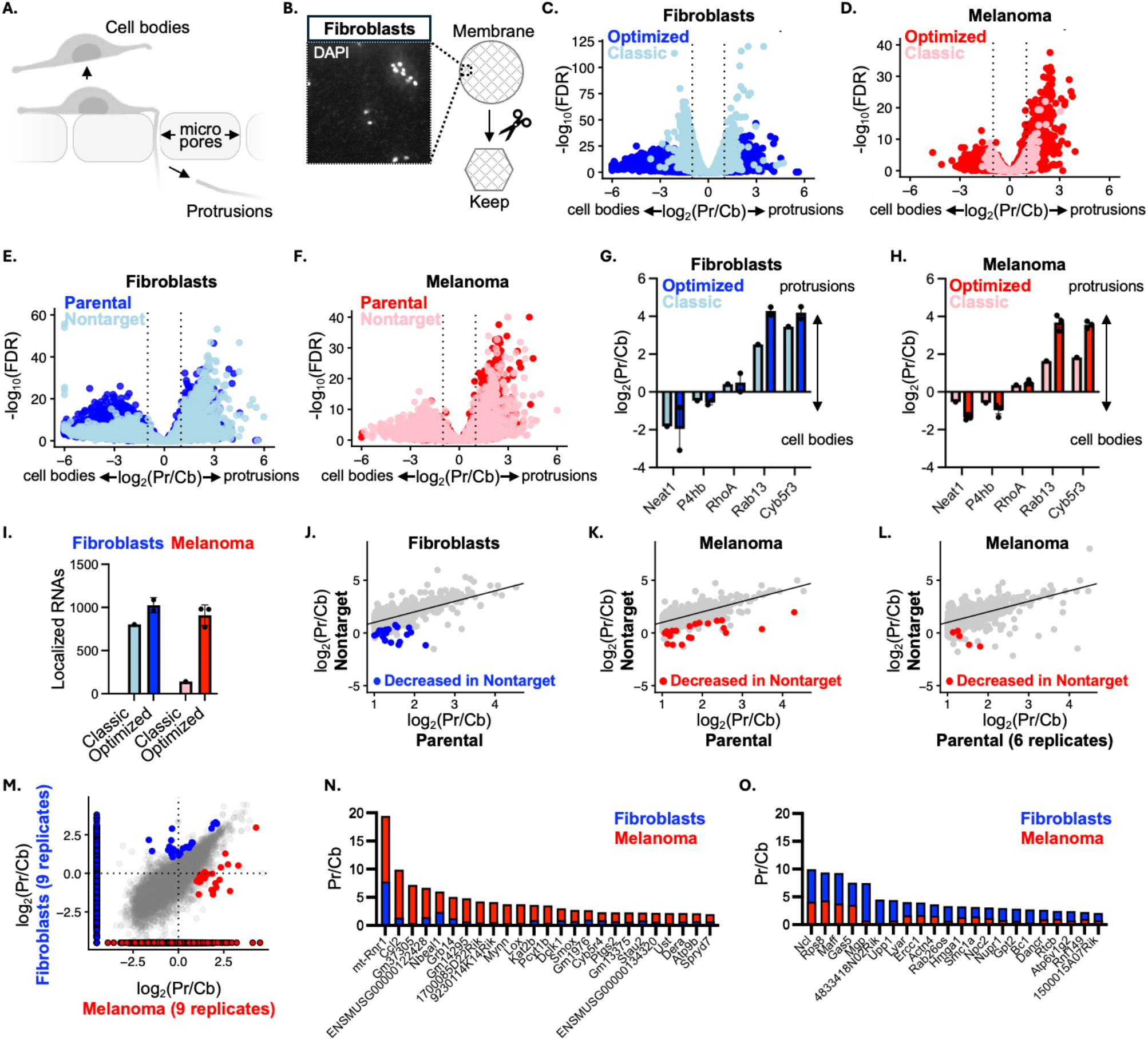
Optimized fractionation improves sensitivity and reproducibility of transcriptome-wide detection of protrusion-localized RNAs. **A.** Schematic of fractionation. Cell bodies are scraped from the top of the membrane and protrusions are lysed from within and beneath the membrane. **B.** Residual nuclei remain at the periphery of the membrane even after washing. The residual nuclei are avoided by cutting out and processing only the center of the membrane. **C-F.** Fractionation RNA-seq comparing the classic versus optimized methods (C-D) or the optimized method before and after single-cell cloning (E-F). **G-H.** RNA localization of representative RNAs from (C-F). Optimized fractionation can distinguish protrusion-localized RNAs (*Rab13* and *Cyb5r3)* from nuclear (*Neat1*) and cell body (*P4hb* and *RhoA*) RNAs. **I.** Total number of localized RNAs detected using log_2_(Pr/Cb) > 1 and FDR < 0.05 threshold. In (G-I), optimized bars combine parental and nontarget data points. **J-L.** Comparison of nontarget and parental data from (E-F). RNAs with significantly decreased localization after single-cell cloning are shown in color. **M.** Comparison of localized RNAs in melanoma versus fibroblast cells. Protrusion-localized RNAs that are significantly more localized in fibroblasts or melanoma cells are blue or red, respectively. RNAs that are expressed only in fibroblasts or melanoma cells are blue and red with a black outline, respectively. In (J-M), significantly changed localization is defined as log_2_(Fold change in localization) > 1 and FDR < 0.1. **N-O.** Fold protrusion enrichment for RNAs that are statistically significantly more localized in melanoma or fibroblastic cells, respectively. All optimized experiments used 3 biological replicates unless otherwise noted.

Several groups have attempted to address these challenges through protrusion stimulation, fixation, and/or extended incubation steps (Dermit et al., 2020; Mardakheh et al., 2015; Mili et al., 2008; Wang et al., 2017), resulting in multiple variations of the classic protocol that vary in both complexity and efficacy.

Here, we present a modified and standardized version of the classic Transwell fractionation method that we optimized for use in non-neuronal cells. The method improves the sensitivity to RNA spatial distribution between protrusions and cell bodies, increases reproducibility across experiments and requires less starting material, all while remaining simple to implement using standard laboratory equipment. By applying our optimized fractionation method to multiple mouse cell lines and genetic mutants, we are able to identify commonalities between RNAs and assign over half of all protrusion-localized RNAs into one of six subsets, which we call “categories”.

Four of the protrusion-localized categories are defined based on known or predicted co-regulation of the RNAs, and we thus refer to them as “co-regulatory categories”. These include APC/KIF1C-dependent RNAs, ribosomal protein mRNAs, other TOP-containing mRNAs and some, but not all, mitochondrially-localized RNAs. Based on research on APC/KIF1C-dependent RNAs, which is the best studied category to date, we predict that RNAs within a category not only share regulatory networks but are also more likely to share downstream molecular outcomes and biological functions. An implication of this prediction is that there may be as many distinct molecular outcomes for protrusion localization of RNAs as there are co-regulatory categories. Our work identifying co-regulatory categories is a first step to testing that hypothesis.

We also define two temporary categories: long non-coding RNAs and pseudogenes. Because these are defined by gene-type instead of shared regulation, they do not meet our definition of co-regulatory categories. However, temporarily grouping them as we have done here draws attention to the peculiarly of their localization, which cannot be explained by current understanding of RNA localization to protrusions. Thus, categorization, even as a placeholder, helps direct attention to conceptual blind spots. Finally, we show that our identified categories also favor protrusions in diverse human non-neuronal cell lines, suggesting that the unifying principles of RNA localization to protrusions, including functional implications, are likely to be conserved.

## RESULTS

### An optimized Transwell fractionation method for non-neuronal cells

We have previously used the classic Transwell method to study RNA localization to protrusions in mouse melanoma cells (Norris & Mendell, 2023). During those studies, we noticed total RNA yield from protrusion fractions was inversely correlated with protrusion-localization signal sensitivity, which suggested residual cell bodies may be contaminating the protrusion fractions. Indeed, DAPI staining confirmed the presence of cell bodies at the edges of membranes even after extensive washing (Figure 1B). To address this, we used a blade to cut out the interior ∼75% of the membrane instead of dislodging and collecting the entire membrane. To further improve standardization, we also excluded all fixation steps or extended incubations, which can add variability across experiments, and used column purification instead of Trizol. Remarkably, these interventions, which we together call “optimized fractionation”, improved the reproducibility and sensitivity of the fractionations, which we describe below.

We applied the optimized fractionation protocol to YUMM1.7 mouse metastatic melanoma cells and NIH3T3 mouse embryonic fibroblasts, two cell types previously studied by ourselves and others using the classic fractionation method (Figure 1C-F)(Norris & Mendell, 2023; Wang et al., 2017). Compared to these classic benchmarks, our optimized method yields >3-fold higher sensitivity, while maintaining specificity to protrusion RNAs. For example, the protrusion-localized mRNA *Rab13* improves from ∼3-5 fold enriched to ∼14-17 fold enriched in protrusions using the optimized method, while the nuclear RNA *Neat1* remains or becomes even more depleted from protrusions (Figure 1G-H). We also detected substantially more localized RNAs, increasing from ∼500 with the classic method to ∼900 with the optimized method, on average per experiment (Figure 1I). Thus, optimized fractionation detects more localized RNAs with more sensitivity than the classic method.

Under our optimized conditions, a single membrane generates ∼25-100 ng total RNA from the protrusion fraction, which has traditionally been the limiting step, and is sufficient for RNA-sequencing. This is a dramatic decrease from the 3-6 membranes and 1 ug RNA per replicate that is standard in the classic protocol. Across 40+ membranes we prepared during optimization, 100% of cell body samples had suitable yield and RNA integrity score (RIN) > 9.4, and 87.5% of protrusion samples had suitable yield and RIN > 8. These results demonstrate that optimized fractionation routinely generates high-quality RNA suitable for RNA-sequencing.

To test reproducibility, we measured RNA localization after single-cell cloning with CRISPR/Cas9 and a non-targeting guide RNA. We chose the single-cell cloning procedure as a test case because it is a commonly used genetic manipulation but is also known to introduce clonal variability, especially with respect to morphology of cell protrusions (Wu et al., 2020). We wanted to see if optimized fractionation was resistant to this technical artifact. Using optimized fractionation, 19 out of 948 localized RNAs in fibroblasts and 21 out of 715 localized RNAs in melanoma cells had decreased protrusion-localization in non-targeted clones compared to parental cells, demonstrating there is a limited amount of interference or “false positives” due to the single-cell cloning procedure (Figure 1J-K). In melanoma cells, the number of false-positive mis-localized RNAs dropped to five when we increased from three to six biological replicates for the parental cells (Figure 1L). Taken together, optimized fractionation maintains high sensitivity and reproducibility even after manipulations known to affect cell protrusions.

### RNA localization to non-neuronal cell protrusions is predominantly cell-type independent

The data above demonstrate that optimized fractionation is a robust and sensitive method capable of detecting protrusion-localized RNAs. Thus, we next wanted to use the method to probe for general principles, such as how RNA localization to protrusions varies across cell types and to what extent localized RNAs use similar or distinct regulatory mechanisms. To this end, we next directly compared our melanoma and fibroblast datasets. Although both cell lines are non-neuronal mesenchymal cells from mice, they are distinct in that the fibroblasts are embryonic and healthy, while the melanoma cells are adult and diseased.

Despite these differences, we find strong correlation between the localization of RNAs in both cell types (Pearson correlation = 0.79)(Figure 1M). 1,249 RNAs localize to protrusions in at least one cell type and half (608) localize to protrusions in both cell types, demonstrating that a substantial number of RNAs localize to protrusions regardless of cell-type identity. The remaining 641 RNAs that do not localize in both cell types fall into three groups. The first group contains 142 RNAs that are expressed in only one of the cell types. The second group contains 44 RNAs that are expressed in both cell types and but have statistically significantly different localization patterns in the two cell types. The last group contains 455 RNAs that are expressed in both cell types but only meet our cutoff for localization in one of the cell types. This final, and largest group, is thus more likely due to cut-off stringency than meaningful biological changes in localization between the cell types. As such, we consider our estimate of 608 RNAs that are localized in both cell-types to be a conservative underestimate.

The substantial similarity in RNA localization between the two cell types is underscored by the fact that, of the RNAs that are expressed in both cell types, only 44 RNAs localize significantly differently between the two cell types, with 20 having increased localization in melanoma, and 24 having increased localization in fibroblasts (Figure 1 M-O). These rare, differentially localized RNAs are thus good candidates for biologically important, cell-type specific processes that are regulated specifically by RNA localization, such as a function in metastatic phenotypes. Taken together, these data demonstrate that RNA localization to protrusions is broadly the same across non-neuronal cells with limited cell type specificity, and that cell-type specific gene expression is more likely to be regulated at the transcription level than at the localization level.

### KIF1C is a broadly acting regulator of RNA localization to protrusions

The robust and high-throughput nature of optimized fractionation now make it feasible to probe the broader regulatory networks underlying RNA localization to protrusions. To this end, we chose to focus on the regulatory role of the kinesin motor protein KIF1C. We chose KIF1C for two reasons. First, KIF1C has previously been shown to traffic seven mRNAs to protrusions, including its own mRNA (Norris & Mendell, 2023; Pichon et al., 2021).

However, as those seven KIF1C-dependent mRNAs were originally discovered using a candidate-based approach, we predicted there may be additional, not yet characterized KIF1C-dependent RNAs that would be detectable using our unbiased method. Second, we wanted to further examine the enigmatic relationship we previously uncovered between *Kif1c* mRNA localization and KIF1C’s role in mRNA trafficking. Specifically, we found that KIF1C protein can properly traffic other KIF1C-dependent mRNAs even when its own mRNA is mis-localized and we wanted to extend that analysis transcriptome-wide (Norris & Mendell, 2023; Pichon et al., 2021).

We applied optimized fractionation to *Kif1c^ΔLOF^* melanoma cells, which lack KIF1C protein, and *Kif1c^Δcis^* cells, which have mis-localized *Kif1c* mRNA due to genetic removal of the *cis-* localization element but still express KIF1C protein (Norris & Mendell, 2023). Unexpectedly, we find 15 RNAs have decreased localization in *Kif1c^Δcis^* cells (Log_2_FC > 1 and FDR < 0.1), suggesting that *Kif1c* mRNA localization does regulate localization of some other RNAs.

Consistent with our previous report, however, only one of the *Kif1c^Δcis^*-affected mRNAs overlaps with the previously defined list of KIF1C-dependent RNAs, which is the *Kif1c* mRNA itself (Figure 2A-B). Taken together, this agrees with our earlier model that *Kif1c* mRNA mis-localization disrupts some functions of KIF1C, while other functions of KIF1C are independent of mRNA localization.

**Figure 2.**
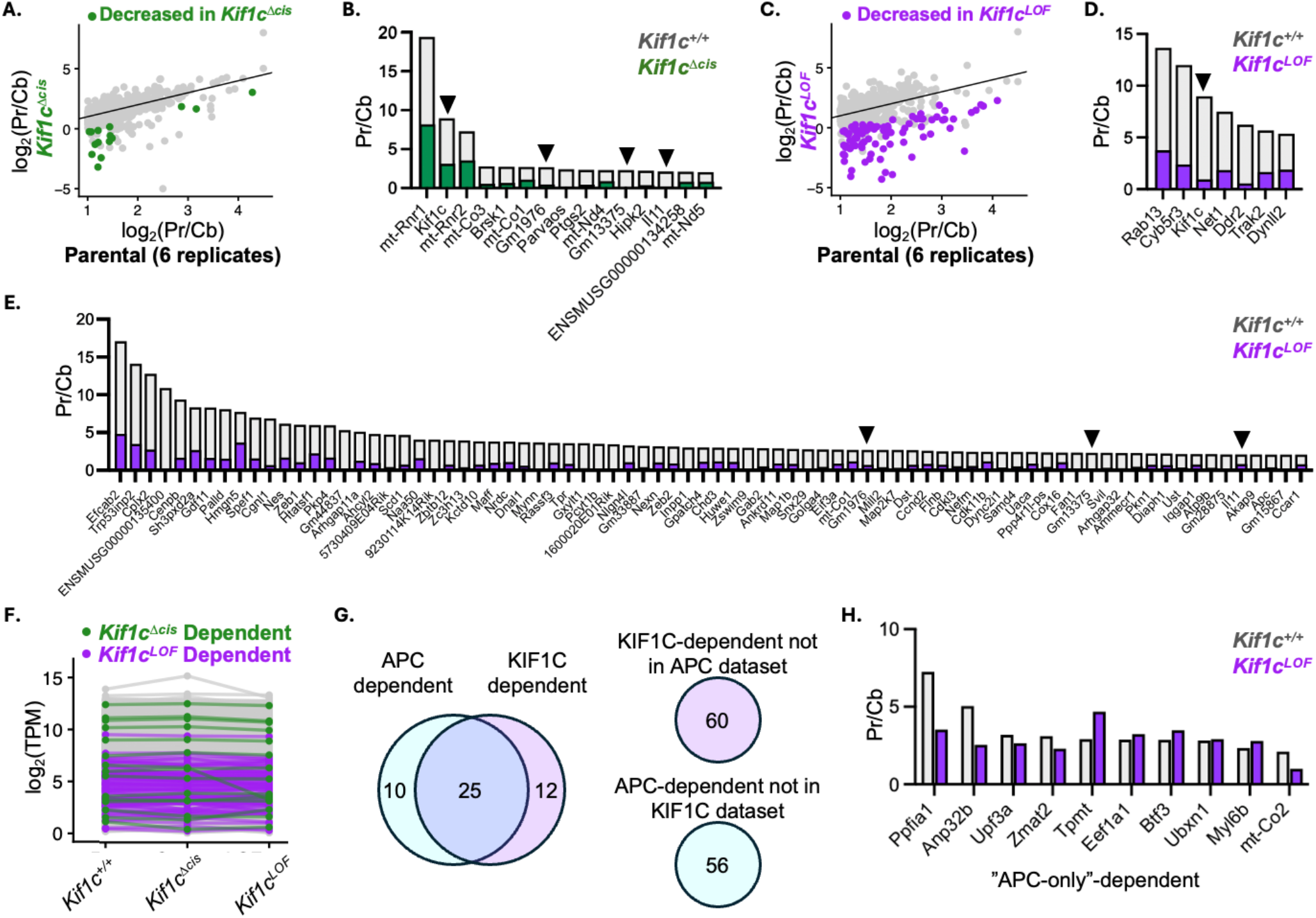
Optimized fractionation identifies 97 KIF1C-dependent RNAs in melanoma cells. **A.** Comparison of localized RNAs in *Kif1c^+/+^*melanoma cells compared to *Kif1c^Δcis^* cells. RNAs with significantly decreased localization in *Kif1c^Δcis^* cells are shown in color. **B.** Fold protrusion enrichment for the 15 RNAs that are significantly less localized in *Kif1c^Δcis^* cells. **C.** Comparison of localized RNAs in *Kif1c^+/+^* melanoma cells compared to *Kif1c^LOF^* cells. RNAs with significantly decreased localization in *Kif1c^LOF^* cells are shown in color. **D-E.** Fold protrusion enrichment for the 7 previously known *Kif1c^LOF^*-dependent RNAs (D.) and 80 newly identified *Kif1c^LOF^*-dependent RNAs (E.). Arrowheads in B, D and E denote RNAs affected in both *Kif1c^Δcis^* and *Kif1c^LOF^* cells. **F.** Average cell body TPM transcriptome-wide plotted across cell lines. **G.** Overlap between KIF1C-dependent and APC-dependent RNAs. **H.** Fold protrusion enrichment for APC-dependent/KIF1C-independent RNAs in *Kif1c^+/+^*and *Kif1c^LOF^* cells. All experiments used 3 biological replicates unless otherwise noted.

In *Kif1c^ΔLOF^* cells we observe substantially more disruption, with a total of 87 mis-localized RNAs (Log_2_FC > 1 and FDR < 0.1)(Figure 2C). This includes all seven previously reported KIF1C-dependent mRNAs, as well as 80 additional RNAs (Figure 2D-E). Notably, the group of KIF1C-dependent RNAs includes ten long non-coding RNAs but still has only four transcripts that overlap with *Kif1c^Δcis^*-affected RNAs (arrowheads in Figure 2B, D and E).

From here forward, we will jointly refer to the 97 total RNAs affected by *Kif1c^Δcis^* and/or *Kif1c^LOF^* as KIF1C-dependent. Although KIF1C-dependent RNAs are less protrusion-localized in *Kif1c^Δcis^* and/or *Kif1c^LOF^* cells, they are still expressed at normal levels in cell bodies (Figure 2F). These data demonstrate that, while KIF1C does act more broadly than previously demonstrated, it is not universally responsible for protrusion-localization and instead modulates localization of a distinct subset of RNAs.

The seven previously known KIF1C-dependent mRNAs also require the tumor-suppressor APC for localization, and we next wanted to determine to what extent this is true for our newly identified KIF1C-dependent RNAs. To this end, we compared our KIF1C-dependent RNAs to a published list of APC-dependent RNAs (Wang et al., 2017). As these gene lists were generated in different cell types (melanoma cells and fibroblasts, respectively) with different methods (optimized versus classic fractionation), we first defined the set of RNAs that were expressed and localized in control cells from both experiments. This identified a set of 35 APC-dependent and 37 KIF1C-dependent RNAs. When we compare these lists, more than half (25/47) are co-regulated by both proteins (Figure 2G). This incomplete overlap could be due to distinct regulatory processes for some of the RNAs or may simply be due to the cutoffs we chose for inclusion in the initial lists, namely a 2-fold or greater change in localization when the *trans-*factor is missing. Manual analysis of the data reveals that, while the cutoff may explain a few genes (especially *Ppfia1, Anp32b* and *mt-Co2*), a strict cutoff does not account for all of the mutually exclusive targets. Indeed, some APC-dependent RNAs are unaffected or even have increased localization in *KIF1C^LOF^* cells, suggesting that these RNAs truly do not require KIF1C for localization (Figure 2H). The observation that KIF1C and APC regulate both common and distinct subsets of RNAs is consistent with a model in which protrusion-localized RNAs are a heterogenous population that can be subset based on shared features, some of which may be overlapping and/or hierarchical.

### Identification of six categories of localized RNAs based on shared features

When viewed together, our melanoma and fibroblast datasets reveal 608 RNAs that are protrusion-localized in both cell types. Based on the work described above and previous results, we can now assign 139 of those RNAs to one of two distinct categories based on their regulatory relationships.

The first category includes any RNA that is KIF1C and/or APC-dependent (from here on called APC/KIF1C-dependent), and accounts for 66 of the 608 protrusion-localized RNAs (purple dots, Figure 3A). This category includes many of the particularly highly localized RNAs. For now, we include the “APC-only” dependent RNAs with the rest of the KIF1C-dependent RNAs but designate them with a black outline in Figure 3A. We expect future research will be able to further subset this category based on the differential activities of the *trans-*factors. The second category of co-regulated localized RNAs is made up of the ribosomal protein mRNAs (blue dots, Figure 3A). Consistent with previous reports, almost all ribosomal protein mRNAs (73/75) localize to protrusions in both cell types, with the two excluded mRNAs barely missing the cutoff.

**Figure 3.**
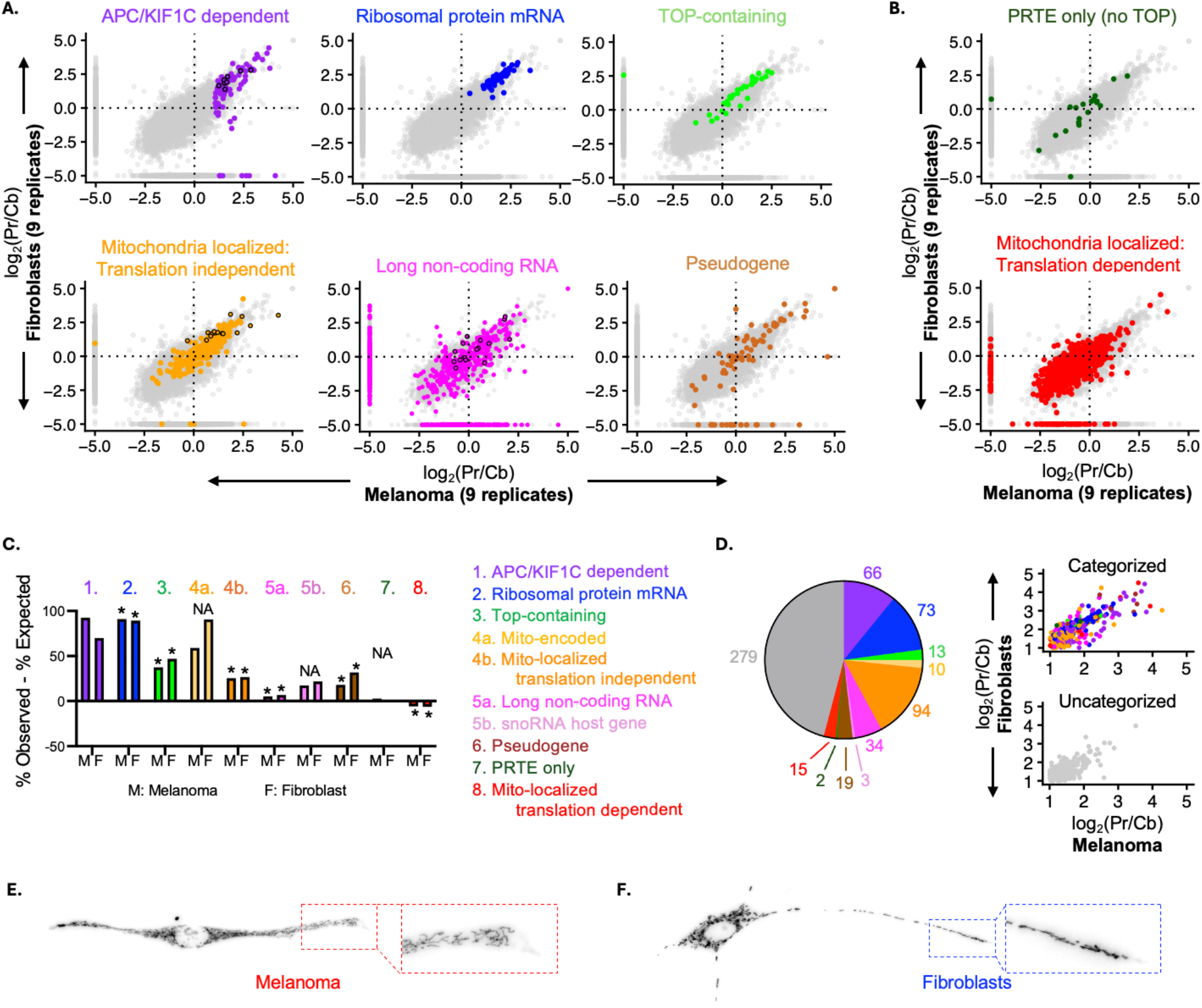
Half of protrusion-localized RNAs fall into one of six categories. **A-B.** Fractionation RNA-seq plots highlighting all RNAs of a given subset/category in melanoma and fibroblastic cells; all other RNAs are shown in gray. The upper right quadrant of each plot represents protrusions from both cell types. **A.** Categories of protrusion-localized RNAs. Dots with black outlines represent subcategories as follows: Purple: “APC-only” dependent. Light orange: Mitochondrially-encoded RNAs. Pink: snoRNA host genes. **B.** 2 subsets of co-regulated RNAs that are not over-represented in protrusions. **C.** Plot showing degree of over-representation of each category. Note that APC/KIF1C-dependent RNAs are a circular argument as they were initially defined by their presence in protrusions. Chi-square test for categories with at least 5 expected RNAs: * p < 0.05. Actual p-values in Supplementary Table 1. **D.** Contribution (left) and localization (right) of each category in the protrusions of melanoma and fibroblast cells. In (C and D) the 10 KIF1C-dependent long non-coding RNAs are only represented in the APC/KIF1C-dependent category. **E-F.** Mitochondria localize to protrusions in melanoma cells (E) and fibroblasts (F).

Based on the presence of these co-regulated categories and evidence that both categories result in distinct molecular outcomes, we propose a model in which subsets of protrusion-localized RNAs use shared regulatory networks to achieve common functional outcomes. This model makes two predictions. First, that there are more, yet unidentified categories of protrusion-localized RNAs. Second, that identification of the missing categories will point the way toward novel functional outcomes.

To identify more potential co-regulatory categories, we screened the remaining 469 uncategorized protrusion-localized RNAs for known or predicted similarities, such as how the RNAs are regulated, their cellular function and/or their gene-type. This analysis revealed the following four categories, which bring the total number of categories to six: #3) other TOP-containing mRNAs (excluding ribosomal protein mRNAs), #4) translation-independent mitochondrially localized RNAs, #5) long non-coding RNAs and #6) pseudogenes. Like ribosomal protein mRNAs, each of these categories are over-represented in protrusions compared to cell bodies (Figure 3C). When combined, the six categories account for more than half of all RNAs that localize in both melanoma cells and fibroblasts (312/608)(Figure 3D). As described more below, the presence of these specific categories, and the absence of similar, related categories, provides insight into which specific RNAs are selected for protrusion localization and underscores that RNA localization to protrusions is not only for protein coding RNAs.

### Ribosomal protein mRNAs and other TOP-containing mRNAs are over-represented in protrusions

All ribosomal protein mRNAs contain pyrimidine-rich 5’TOP motifs, which are believed to regulate their subcellular localization. However, the exact nature of 5’TOP motifs and their annotation throughout the transcriptome is still being perfected and refined. Indeed, multiple other pyrimidine-rich “TOP-like” motifs have been reported such as the “*cis*-element upstream of the initiation codon” (CUIC) and “pyrimidine rich translation element” (PRTE) (Hsieh et al., 2012; Shigeoka et al., 2019). To what extent these motifs are redundant or distinct is yet to be determined, and individual transcripts can contain multiple motifs. For these reasons, we separate ribosomal protein mRNAs into their own category, as they share a similar biological function (ribosomal proteins) in addition to the presence of a 5’TOP. After removing ribosomal protein mRNAs, our dataset contains 31 other mRNAs that are annotated as containing a 5’TOP motif and constitute our third category (light green dots in Figure 3A). Strikingly, we do not see strong protrusion localization for mRNAs that contain only the “TOP-like” PRTE-motif (dark green dots in Figure 3B). These data imply that “TOP” and “TOP-like” motifs are not equivalent in regard to their role in subcellular localization in non-neuronal cells.

### Translation-independent, but not translation-dependent, mitochondrially-localized RNAs localize to protrusions

Mitochondria, due to their origination as symbionts, have their own genome consisting of 13 protein coding genes. We find that the mRNA for these mitochondrially-encoded genes are robustly localized to protrusions (orange dots with black outlines in Figure 3A). This may be explained by the fact that mitochondria themselves localize to protrusions in fibroblasts and melanoma cells, as localization of the whole organelle would result in localization of the mitochondrially encoded transcripts (Figure 3E-F), assuming that the localized mitochondria retain their mitochondrial DNA.

In addition to mitochondrially-encoded RNAs, recent research has cataloged around 1,500 nuclear-encoded RNAs that localize to the outer membrane of mitochondria under certain conditions (from here on referred to as mito-localized RNAs)(Fazal et al., 2019; Luo et al., 2025). These nuclear-encoded mito-localized RNAs fall into two broad categories: those that localize co-translationally and are translation-dependent, and those use *cis-*elements within the RNA sequence and are translation-independent. Intriguingly, these mechanistically distinct subsets appear separately in our fractionated datasets. While translation-independent mito-localized RNAs are statistically over-represented in protrusions in both fibroblasts and melanoma cells, translation-dependent mito-localized RNAs are statistically under-represented (compare orange dots in 3A and red dots in 3B). These data suggest that RNAs not only localize to mitochondria through distinct regulatory mechanisms (translation dependent vs independent) but may also localize to spatially distinct mitochondria. Alternatively, translation-independent mito-localized RNAs may reach protrusions in a mitochondria independent way. Thus, our results uncover an unexpected spatial element to mitochondrial localized RNAs that segregates alongside previously established regulatory mechanisms. Notably, this spatial segregation was not detected or predicted by the RNA localization methods originally used to define mitochondrial RNA localization mechanisms, and the implications to mitochondrial and/or cellular biology will require further analysis.

### Long non-coding RNAs and pseudogenes are over-represented in protrusions

Unlike the categories discussed so far, in which we have grouped RNAs by their co-regulatory relationships, we define the last two categories, long non-coding RNAs and pseudogenes, by their gene types. For both categories, the over-representation in protrusions is striking as there is no intuitive reason for them to be enriched there.

Pseudogenes, long thought to be relics, are increasingly reported to have cellular functions, and their localization to protrusions could suggest a functional role for the localized pseudogenes in particular. Likewise, although there are now multiple examples of long non-coding RNAs with function outside the nucleus, the utility of their localization specifically to protrusions is unclear. We note that snoRNA host genes in particular appear to favor protrusions (pink dots with black outlines in Figure 3A). While the main function of these host genes is believed to be generation of snoRNAs, which function in the nucleolus, the presence of the host transcript in protrusions suggests an additional regulatory and/or functional use for these transcripts and is consistent with recent reports of secondary functions for some snoRNA host genes (Chen et al., 2023). In all cases, the purpose of localizing large numbers of long non-coding RNAs and/or pseudogenes specifically to protrusions reveals a layer of post-transcriptional regulation that is yet to be understood.

### The categories of protrusion-localized RNAs are conserved in humans

Our analysis above reveals that over half of protrusion-localized RNAs in mouse non-neuronal cells can be subset into categories based on shared features, including potential or established co-regulatory mechanisms. To determine if these categories represent broadly acting unifying principles, we next wanted to see if and to what extent they are present in other cell types, including human cell types. To this end, we expanded our analysis to include six human non-neuronal cell lines, including MDAMB231 human breast cancer cells, which are one of the most comprehensively studied human non-neuronal mesenchymal cell lines in regard to RNA localization to protrusions (Mardakheh et al., 2015; Moriarty et al., 2022). When we analyzed published fractionation data for the panel of human cell lines, a modest but clear preference for protrusions is seen for all six categories, although the quantitative strength of these trends is weaker (Figure 4C). For example, while the majority of the transcripts in each category prefer protrusions to cell bodies (log_2_(Pr/Cb) > 0), relatively few are above our strict cutoff for protrusions (log_2_(Pr/Cb) > 1). For this reason, we applied a more modest cutoff of log_2_(Pr/Cb) > 0.5 for the human dataset when quantifying the amount of RNAs localized in each category (Figure 4D). Whether the weaker localization values reflect true biological differences or are an artifact of the different fractionation methods is not clear. Regardless, the trends are sufficiently strong to conclude that these categories of RNAs represent conserved organizational strategies and imply that RNA localization is biologically useful in each of these cell types.

**Figure 4.**
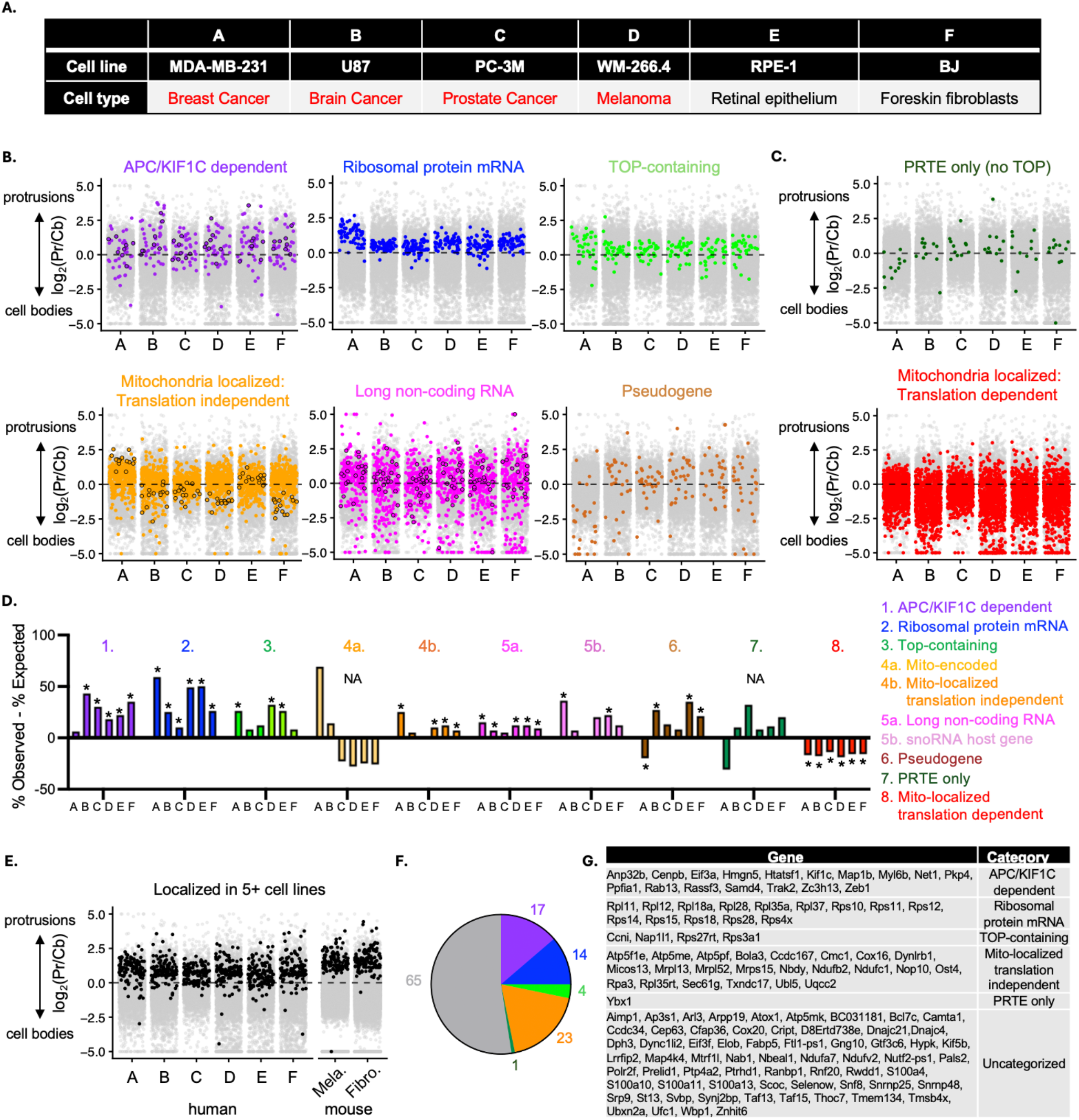
Categories of protrusion-localized RNAs are modular and conserved in humans. **A.** List of cell lines plotted in (B-E). Lines listed in red are cancerous. **B-C.** RNA-seq fractionation plots highlighting all RNAs of a given subset/category; all other RNAs are shown in gray. **B.** Categories of protrusion-localized RNAs originally defined in mice. Dots with black outlines represent subcategories as follows: Purple: “APC-only” dependent. Light orange: Mitochondrially-encoded RNAs. Pink: snoRNA host genes. **C.** 2 additional subsets of co-regulated RNAs. **D.** Plot showing degree of over-representation of each category. Chi-square test for categories with at least 5 expected RNAs: * p < 0.05. Actual p-values in Supplementary Table 2. **E.** Plot showing RNAs that localize to protrusions in at least four human cell lines and one mouse cell line. **F.** Contribution of each category to the conserved localized RNAs highlighted in (E). **F.** Conserved protrusion-localized RNAs and their respective categories.

### Co-regulatory categories of RNAs show cell type specific modularity

The comparison of many cell lines not only allowed us to identify conserved similarities but also revealed multiple striking exceptions. One example is the behavior of “mitochondrially-related” RNAs. In mouse cells we found that mitochondrially-encoded RNAs and mito-localized translation independent RNAs both localized to protrusions, while mito-localized translation dependent RNAs did not. While this pattern is also true for two of the six human cell lines, in the other four human cell lines, mitochondrially-encoded RNAs are not protrusion-localized. Thus, there are at least three distinct groups of RNAs that, while all affiliated with mitochondria, can be distinguished within our spatial datasets. Furthermore, localization of each group to protrusions, or not, appears to be modular, as localization of one group can change without affecting localization of the other groups.

This modularity of localization is also apparent when considering pseudogenes and PRTE-containing RNAs. PRTE-motifs, which are similar to but distinct from TOP-motifs, do not drive protrusion localization in mouse non-neuronal cells (dark green dots in Figure 3B).

However, in five out of six human non-neuronal cell lines, PRTE-containing mRNAs do localize to protrusions, in many cases as well or better than TOP-containing mRNAs, with human breast cancer cells being the sole exception (dark green dots in Figure 4C).

Importantly, in human breast cancer cells TOP-containing mRNAs do localize to protrusions, while PRTE-containing RNAs do not. This suggests that, like mito-localized RNAs, PRTE-containing RNAs can be modularly regulated, and that changes to one category (e.g. PRTE-containing) does not affect other categories (e.g. TOP-containing RNAs). We conclude that PRTE-containing RNAs represent a 7^th^, distinct co-regulatory category of localized RNAs that is modularly de-localized in the two mouse non-neuronal cell lines we have studied so far.

A third example of modularity is the lack of localization for both the APC/KIF1C dependent and pseudogene categories specifically in human breast cancer cells. Like PRTE-containing RNAs, APC/KIF1C dependent and pseudogene localization is de-localized in just this cell line. It is intriguing to speculate that this cell-type specific change could be a biologically relevant dysregulation. Indeed, this human breast cancer cell line has been reported to have less mechanoactivity during cell migration, which correlates with altered RNA localization behavior (Moriarty et al., 2022). It also suggests that pseudogenes may in fact have shared underlying co-regulation, and may be a bona fide “co-regulatory” category. Future research will be able to test this hypothesis and determine to what extent pseudogene mis-localization translates into cell biological outcomes. Taken together, these data suggest that localization of RNAs is a modular process, and cells can pick and choose which categories to localize depending on the needs of that cell type.

### A core set of conserved, protrusion-localized RNAs

So far we have focused on RNA localization at the level of categories. We next wanted to ask which specific localized transcripts are conserved across diverse cell types and species. To this end, we looked for RNAs that are localized to protrusions in at least 4 of the human cell lines and at least one of the mouse cell lines (log_2_(Pr/Cb) > 0.5 for human or > 1 for mice; FDR < 0.05). We identify 124 RNAs that meet our criteria, nearly half of which (59/124) fall into one of our six categories (Figure 4E-G). We propose that these conserved localized RNAs are particularly good representatives for understanding the role and regulation of RNA localization in cell biology and physiology, and that lessons learned from a single one of these RNAs are likely to extend to other RNAs within the same category.

## DISCUSSION

Here, we present an optimized version of the classic Transwell fractionation method for use in non-neuronal cells. The method is more robust, more efficient and more sensitive than the classic technique. By applying optimized fractionation to multiple cell types and genetic mutants, we uncover six distinct categories of localized transcripts, which together account for over half of all protrusion-localized RNAs in mouse non-neuronal mesenchymal cells.

These six categories, as well as at least one additional category, also localize to protrusions in human non-neuronal cells. We propose that localization of each category serves a distinct functional purpose, and that individual cell types can modularly choose which categories to localize or not. Furthermore, the presence of multiple categories, including an unknown number that have not yet been defined, suggests there may be many not yet discovered non-canonical functions for RNA localization to cell protrusions.

Although we find that, in general, RNA localization to protrusions is cell type agnostic, we identify a small collection of cell type specific localized RNAs. The presence of these rare events suggests that localization of these RNAs in particular may have cell type specific functions and could be an informative route for future analysis. Optimized fractionation, which can be readily adapted to diverse cell types and manipulations, provides a robust means for dissecting these mechanisms in the future.

The comparison of RNA localization to protrusions in neuronal versus non-neuronal cells leads to an intriguing paradox. On one hand, there is a clear functional disconnect between the two cell types: while canonical mRNA localization leads to localized proteins in neuronal cells, this relationship is absent in non-neuronal cells. On the other hand, many of the key regulators appear to be shared in neurons and non-neuronal cells. For example, KIF1C, APC, LARP1, GA-rich elements and TOP motifs are all believed to regulate RNA localization in neurons and non-neuronal cells alike (Arora et al., 2022; Goering et al., 2023; Loedige et al., 2023; Nagel et al., 2023; Preitner et al., 2014). Thus, different types of cells have evolved to use common machinery to achieve apparently distinct molecular outcomes. This is consistent with studies in other cell types that likewise concluded that RNA localization is a general mechanism of spatial gene regulation, rather than a cell type-specific phenomenon (Goering et al., 2023). Future work will have to tease out where the similarities end and how the unique functional outcomes in each cell type arise.

Our model that protrusion-localized RNAs use shared regulatory networks to achieve distinct functional outcomes suggests that identification of subsets of RNAs will help uncover and define novel molecular mechanisms. Here, we have begun the process of systematically cataloging localized RNAs by defining seven categories. Although these categories are preliminary, and two are defined by gene type as opposed to shared regulation, the results nonetheless point toward open questions and important next steps.

We, like others, report the localization of virtually all ribosomal protein mRNAs to protrusions. Although this arrangement has been repeatedly described, the functional utility of this localization remains unclear, especially given that ribosomes are constructed within the nucleolus, far from protrusions. The fact that LARP1 contributes to mRNA stability as well as mRNA localization has made it difficult to mechanistically tease out regulatory processes. However, optimized fractionation, which can detect even subtle regulatory changes, now simplifies the rapid characterization of *trans*-factors and lends itself to whatever perturbations may be necessary for addressing this enigmatic phenomenon.

We find that TOP-containing transcripts, with and without an additional PRTE, are over-represented in non-neuronal cell protrusions, while transcripts with only a PRTE are more evenly distributed or absent from protrusions in both mouse melanoma and fibroblasts. This stratification of localization behavior is distinct from previous work in epithelial and neuronal cells, in which both the TOP and PRTE motifs could drive basal and neurite localization (Goering et al., 2023). Intriguingly, PRTE-containing transcripts do localize to protrusions in some, but not all, human cell lines that we examined. This is consistent with the idea that regulatory principles of RNA localization are modular and may or may not be deployed in the same way across diverse cell types, which may in turn help explain divergent functional outcomes.

We see a similar unanticipated regulatory hierarchy with mitochondrial-localized RNAs. Prior studies using proximity ligation have identified two sets of nuclear-encoded, mitochondrially localized RNAs: those that require translation for localization, and those that are translation independent. While the proximity studies treated all mitochondria as a single pool, our work provides an additional dimension of analysis by separating the mitochondria based on their subcellular localization and reveals that only translation-independent mitochondrially localized RNAs reach protrusions. Surprisingly, mitochondrially-encoded RNAs also show modularly controlled protrusion-localization, as they localize to protrusions in half of all examined cell lines. Whether these observations are due to heterogeneity amongst the mitochondria, the RNA, or both will require future analysis.

We observe strong enrichment for both pseudogenes and long non-coding RNAs in non-neuronal cell protrusions. While we consider it unlikely that the long non-coding RNAs and pseudogenes are all co-regulated, we have currently categorized them this way due to a lack of better information about their regulation. However, future work will provide clarity in this area. Indeed, here we report ten long-noncoding RNAs that require the KIF1C for localization, revealing that long non-coding RNAs and mRNAs can be co-regulated.

While Transwell fractionation is powerful for its ability to provide high-throughput transcriptome-wide analysis, it comes at the cost of detailed spatial information, as all shapes, sizes and types of protrusions are collected indiscriminately. This is an important caveat, as there is evidence that there is diversity to when, where and which RNAs localize within different types of protrusions. The best examples of this are the preference of APC/KIF1C-dependent RNAs for long, microtubule-rich protrusions, and the preference of ribosomal protein mRNAs for smaller, actin-rich protrusions (Wang et al., 2017). However, even with these limitations, we show that optimized fractionation is able to robustly detect specific localization changes upon loss of essential factors. Furthermore, given that our understanding of protrusion-specific preferences is still underdeveloped, the agnosticism of fractionation toward the protrusion subtype makes it a powerful technique that works even without prior knowledge of which type of protrusion to focus on.

In conclusion, we have developed an optimized fractionation method that enables sensitive and reproducible detection of protrusion-localized RNAs in non-neuronal mesenchymal cells. The protocol is compatible with a wide range of experimental manipulations and, because it yields high-quality RNA from a limited number of cells, simplifies generating larger numbers of biological replicates, thereby increasing statistical power in RNA-sequencing experiments. We have applied the optimized fractionation to mouse non-neuronal cells and mutant cell lines and identified multiple categories of conserved, localized RNAs, which we predict share regulatory and functional features. We anticipate our conceptual and methodological approaches will be broadly applicable across diverse cell types, culture conditions, and genetic perturbations, and will help place the enigmatic process of RNA localization to non-neuronal cell protrusions within the greater context of cell biology, physiology and disease.

## MATERIAL AND METHODS

### Cell culture

Established YUMM1.7 and NIH3T3 cell lines were obtained from ATCC and tested negative for mycoplasma. The YUMM1.7 cell line was authenticated after receipt using allele-specific primers. YUMM1.7 cells were cultured in DMEM/F12, HEPES with L-glutamine (GIBCO 11330057) supplemented with 10% FBS (Thermofisher A5209402), 5% nonessential amino acids, and 5% penicillin/streptomycin (GIBCO 11140050 and 15140122). NIH3T3 cells were cultured in DMEM with L-Glutamine (GIBCO 11995073) supplemented with 10% FBS and 5% antibiotic/antimycotic (GIBCO 15240062).

### Optimized fractionation and RNA extraction

Cells were grown to 70-80% confluency. PET Millicell hanging cell culture inserts with 1-um pores sized for six-well plates were coated on the bottom side of the membrane with bovine plasma fibronectin in a sterile hood (Millipore Sigma PTRP06H48 and F1141). Briefly, fibronectin was diluted to 30 µg/mL in sterile DPBS. 500 µl diluted fibronectin was dotted onto the inside of an upturned lid of a 6 well plate. One insert was placed on top of the fibronectin dot such that the bottom side of the membrane was in full contact with the fibronectin, then incubated for 15 minutes at room temperature. Residual fibronectin was aspirated, and inserts were turned upside down and allowed to air-dry membrane facing up for at least 15 minutes at room temperature. Coated and dried inserts were placed in 6 well plates containing 3 mL of complete media per well. Cells were prepared by gently pelleting passaged cells and resuspending in complete media to a final concentration of 4-5×10^5^ cells/mL. 2 mL of cells were added inside each insert such that there was 8-10×10^5^ total cells evenly distributed across each insert. Cells were incubated overnight in a standard cell culture incubator. After 16-24 hours, one plate at a time was brought to a standard lab bench for fractionation. Working one insert at a time, media from inside the insert was aspirated, cells were gently rinsed with 200 ul RNase-free PBS, and the PBS was aspirated. Cell bodies were dislodged from the top of the membrane by scraping inside the insert using a cell scraper (Fisher 08-100-241). Visible clumps of cell bodies were collected into a tube of 600 ul RLT buffer from the Qiagen RNeasy kit (Qiagen 74106) and vortexed 10 seconds. The inside of the scraped insert was swabbed with a cotton swab to remove visible residual cell bodies, then rinsed with 200 ul RNase free PBS. The PBS was aspirated, and the top of the membrane was cleaned by swabbing in all directions with at least three cotton swabs followed by another PBS rinse and aspiration. This cleaning procedure was repeated twice more. The cleaned membrane was cut from the insert using a sharp blade, leaving about 1/8^th^ of an inch at the edges. The cleaned, cut membrane which contained protrusions was placed in a new tube with 600 ul RLT buffer and vortexed for 10 seconds. Isolated fractions were then kept at room temperature in RLT buffer while all additional membranes were fractionated. Once all samples were collected into RLT buffer, all tubes were vortexed for an additional 10 seconds, then the standard protocol for RNeasy was followed, including DNase digestion (Qiagen 79256). RNA was eluted with 30 ul of RNase-free water. The expected concentration of cell bodies depends on how much of the visible cell body clump was added to the tube but is generally between 300 ng/µl and 1 µg/µl. The concentration of protrusions is generally 1-3 ng/µl but can be as low at 0.5 ng/µl. Concentrations above 7 ng/µl generally suggest cell body contamination and result in lower levels of observed protrusion localization. We did no other “pre-screening” of the samples prior to standard quality control for RNA-sequencing.

### RNA-sequencing

Each RNA-seq replicate came from a single membrane. Specifically, each membrane gave one cell body sample and one protrusion sample, which became two paired RNA-seq samples. Below, we refer to these paired cell body-protrusion samples as a single RNA-seq replicate, such that three “replicates” actually contains six samples. During analysis, the protrusions of a given membrane were compared to the cell bodies of the same membrane. 2 ul of RNA for each sample was used for Tapestation. Libraries were generated from total RNA with NEBNext Ultra II Directional Library Prep kit (input 10ng – 1ug total RNA). For all RNA-seq experiments three replicates were used per cell line. The experiments were carried out as follows: YUMM_Parental_experiment_1 was run on the Illumina NextSeq P2, single end x 100 cycles. YUMM_Parental_experiment_2, YUMM_Nontarget, YUMM_*Kif1c^Δcis^,* YUMM_*Kif1c^LOF^*, NIH3T3_Parental and NIH3T3_Nontarget were all carried out on an Illumina Novaseq X Plus, paired end x 150 cycles.

### RNA-seq data analysis

Quality assessment of the reads was done using FastQC (ver 0.11.9) and Trim_galore (ver 0.6.7)(Babraham Institute). Reads were quasi-mapped to the Gencode mouse transcripts (GRCm39 release M36) or human transcripts (GRCh38, release 49) using Salmon (ver 1.7.0)(Patro et al., 2017). Transcript abundance was imported into R (version 4.4.2) with the ‘tximport’ package (ver 1.34.0)(Soneson et al., 2015) and differential expression analysis was carried out using DESeq2 (ver 1.46.0)(Love et al., 2014). Protrusion enrichment values were calculated as protrusion/cell body (PR/CB) using the Wald test within DESeq2 for a given genotype. Differential protrusion enrichments among genotypes were calculated as (protrusion_parental/cellbody_parental)/(protrusion_mutant/cellbody_mutant) using the likelihood ratio test (LRT) within DESeq2. In the plots with gene expression, gene expression was approximated by averaging the cell body TPM values across replicates for each cell line.

### Quantification of localization

Only RNAs with cell body expression of TPM > 1 were considered. RNAs were considered “localized” when Log_2_(Pr/CB) > 1 in mice or Log_2_(Pr/CB) > 0.05 in humans and FDR < 0.05. For the volcano plots, YUMM_Parental_experiment_1 was used for melanoma. For the localization bar plots for *Neat1, P4hb, RhoA*, *Rab13* and *Cyb543*, and the bar plots for total number of localized RNAs: the optimized fibroblast bars show results from NIH3T3_Parental and NIH3T3_Nontarget and the optimized melanoma bars show results from YUMM_Parental_experiment_1, YUMM_Parental_experiment_2, and YUMM_Nontarget. YUMM_Parental_experiment_2 was included in experiments that state 6 replicates were used for melanoma. For identification of significantly affected localized RNAs, localized RNAs were first identified in the parental cell line, then localization values for those RNAs were compared to the localization values in the other cell line of interest.

RNAs were considered significantly decreased if Log_2_((protrusion_parental/cellbody_parental)/(protrusion_mutant/cellbody_mutant)) > 1 and FDR < 0.1. To identify RNAs affected by single cell cloning, NIH3T3_Parental was compared to NIH3T3_Nontarget, and YUMM_Parental_experiment_1 was compared to YUMM_Nontarget. When six replicates were used, YUMM_Parental_experiment_1 and YUMM_Parental_experiment_2 were combined before comparison to YUMM_Nontarget. To identify RNAs changed in *Kif1c^Δcis^* or *Kif1c^LOF^* cell lines, we compared those cell lines to all YUMM1.7 *Kif1c^+/+^* cells, which included the six replicates of YUMM_parental (YUMM_Parental_experiment_1, YUMM_Parental_experiment_2) and the 3 replicates of YUMM_Nontarget. Fold protrusion localization is calculated as 2^Log_2_(Pr/Cb).

NIH3T3_Parental, NIH3T3_Nontarget and NIH3T3_Nontarget2 were combined for experiments using 9 replicates. YUMM_Parental_experiment_1, YUMM_Parental_experiment_2, and YUMM_Nontarget were combined for experiments using 9 replicates. A list of 608 localized RNAs was defined as RNAs with CB TPM >= 1, log_2_(Pr/Cb) > 1 and FDR < 0.05 in both melanoma and fibroblast datasets.

### Comparing classic and optimized protocols

Classic: Data from Wang et al was downloaded from GEO and included four replicates of fractionated NIH3T3 cells. Data from Norris and Mendell was processed using our RNA seq analysis pipeline as described elsewhere in the methods and included two replicates of fractionated YUMM1.7 cells. Optimized: Unpublished data was processed using our RNA seq analysis pipeline as described elsewhere in the methods and included three replicates for each of the following fractionated cell lines: NIH3T3_Parental, NIH3T3_Nontarget, YUMM_Parental_experiment_1, YUMM_Parental_experiment_2, YUMM_Nontarget. The following was applied for all of the datasets: A cutoff of TPM>=1 was applied to cell body fractions. Localized RNAs were defined as log_2_(Pr/Cb)>1 and FDR <0.05.

### Gene category annotations

APC/KIF1C-dependent RNAs were defined as the 97 RNAs mis-localized in *Kif1c^LOF^* and/or *Kif1c*^Δcis^ cells and the 10 “APC-only” RNAs, as described in Figure 2G. Mouse ribosomal protein mRNAs were taken from the Ribosomal Protein Gene Database http://ribosome.med.miyazaki-u.ac.jp/rpg.cgi?mode=orglist&org=Mus%20musculus. Human ribosomal protein mRNAs were taken from the dataset annotated in (Dermit et al., 2020).

TOP-containing and PRTE-only RNAs were defined based on (Hsieh et al., 2012). Mitochondrial encoded RNAs were defined as 13 mitochondrial mRNAs and two mitochondrial ribosomal RNAs. Translation dependent, mitochondrially-localized RNAs were defined as the union between “Short-CDS” RNAs (Luo et al., 2025) and “RNA-dependent” RNAs (Fazal et al., 2019). Translation independent, mitochondrially-localized RNAs were defined as the union between “Long-CDS” RNAs (Luo et al., 2025) and “Ribosome-dependent” RNAs (Fazal et al., 2019). Long non-coding RNAs and pseudogenes were defined by their annotation in GENCODE. snoRNA host genes were defined as genes with SNHG in their name, as well as GAS5 and DANCR. Mouse and Human orthologs were downloaded from Biomart.

### Live cell staining of mitochondria and imaging

250 uL of 1.5 x 10^4^ YUMM cell/mL or NIH3T3 cell/mL were plated in a chambered coverslip coated with 30 µg/mL fibronectin and incubated for 18 hours. Cell media was exchanged for 250 µL 100 nM BioTracker 488 Green Mitochondria Dye in YUMM or 3T3 complete media and incubated at 37°C for 15 minutes. Dye solution was removed and washed with 250 µL sterile PBS before imaging in PBS. Cells were imaged 10 minutes after staining on a Keyence BZ-X710 microscope. Image settings were as follows: 100% LASER power, standard resolution (8-bit), low photobleach, 60X water immersion lens, GFP filter cube (BZ-X Filter GFP – OP-87763). YUMM cells were imaged at a 1/3 s exposure while NIH3T3 cells were imaged at a 1/5 s exposure.

## ACKNOWLEDGEMENTS

We would like to acknowledge Dr. Brandon Le and the IIGB Bioinformatics Core for developing the pipeline to process the RNA-seq data. We thank Dr. Adam Norris for helpful comments on the manuscript and the University of California, Riverside School of Medicine Research Core for their assistance in fluorescence-activated cell sorting. We also thank the Genomics Cores at the University of California Riverside and Los Angeles campuses for high-throughput sequencing.

## Author contributions

BH: Performed experiments and developed methodology. SK, DM and VV: Performed experiments. MLN: Conceptualized the project, performed experiments, developed methodology, analyzed results and wrote the manuscript.

## DATA AVAILABILITY

Data has been deposited to GEO and will be available upon publication.

---

**Supplemental Table 1.** P-values for Chi-square test shown in Figure 3C. NA is shown for categories with < 5 expected observations, for which the Chi-square test is invalid.

|  | 1 | 2 | 3 | 4a | 4b | 5a | 5b | 6 | 7 | 8 |
| --- | --- | --- | --- | --- | --- | --- | --- | --- | --- | --- |
| Melanoma | NA | <0.0001 | <0.0001 | NA | <0.0001 | <0.0001 | NA | <0.0001 | NA | <0.0001 |
| Fibroblast | NA | <0.0001 | NA | NA | <0.0001 | <0.0001 | NA | <0.0001 | NA | <0.0001 |

**Supplemental Table 2.** P-values for Chi-square test shown in Figure 4D. NA is shown for categories with < 5 expected observations, for which the Chi-square test is invalid.

|  | 1 | 2 | 3 | 4a | 4b | 5a | 5b | 6 | 7 | 8 |
| --- | --- | --- | --- | --- | --- | --- | --- | --- | --- | --- |
| A | 0.3364 | <0.0001 | 0.001 | NA | <0.0001 | <0.0001 | 0.0012 | 0.0152 | NA | <0.0001 |
| B | <0.0001 | <0.0001 | 0.246 | NA | 0.0523 | 0.0121 | 0.6291 | <0.0005 | NA | <0.0001 |
| C | <0.0001 | 0.0463 | 0.1074 | NA | 0.7858 | 0.0633 | 1 | 0.0654 | NA | <0.0001 |
| D | 0.0088 | <0.0001 | <0.0001 | NA | <0.0001 | <0.0001 | 0.0533 | 0.4028 | NA | <0.0001 |
| E | 0.0008 | <0.0001 | 0.0005 | NA | <0.0001 | <0.0001 | 0.0104 | <0.0001 | NA | <0.0001 |
| F | <0.0001 | <0.0001 | 0.2460 | NA | 0.0043 | 0.0019 | 0.1243 | 0.0088 | NA | <0.0001 |

